# Female rats consume and prefer oxycodone more than males in a chronic two-bottle oral voluntary choice paradigm

**DOI:** 10.1101/690735

**Authors:** Giulia Zanni, Matthew J. DeSalle, Hannah M. Deutsch, Gordon A. Barr, Amelia J. Eisch

## Abstract

The increased abuse of opioids - such as oxycodone - poses major challenges for health and socioeconomic systems. Human prescription opioid abuse is marked by continuous, voluntary, oral intake, and sex differences. Therefore the field would benefit from a preclinical in-depth characterization of sex differences in a chronic oral voluntary, free choice, and continuous access paradigm. Here we show in an oral oxycodone continuous access two-bottle choice paradigm sex-dependent voluntary drug intake, dependence, and motivation to take the drug. Adult female and male Long-Evans rats were given unlimited, continuous home cage access to two bottles containing water (Control) or one bottle of water and one bottle of oxycodone dissolved in water (Experimental). Most experimental rats voluntarily drank oxycodone (∼10 mg/kg/day) and escalated their intake over 22 weeks. Females self-administered twice as much oxycodone as males, leading to greater blood levels of oxycodone, and engaged in more gnawing behavior. Precipitated withdrawal revealed high levels of dependence in both sexes. Reflecting motivation to drink oxycodone, ascending concentration of citric acid suppressed the intake of oxycodone (Experimental) and the intake of water (Control); however Experimental rats returned to pre-citric acid preference levels whereas Controls rats did not. Thus, female rats consumed and preferred oxycodone more than males in this chronic two-bottle oral choice paradigm. Both sexes displayed many features of human oxycodone abuse, and behavioral pre-screening predicted parameters of intake and withdrawal. This model provides an additional paradigm for understanding mechanisms that mediate long-term voluntary drug use and for exploring potential treatment options.

**HIGHLIGHTS:** Adult rats offered continuous choice of oral oxycodone vs. water preferred oxycodone

Rats self-titrated oxycodone, yet females preferred and escalated more than males

Both sexes were motivated to drink oxycodone, as shown by a citric acid aversion test

Both sexes became dependent on oxycodone, as shown by precipitated withdrawal

Behavioral prescreening predicted later aspects of oxycodone intake and dependence

## 1. INTRODUCTION

Opioid misuse and abuse in the US and globally pose major challenges for health and socioeconomic systems worldwide (Cicero and Ellis, 2017a; Compton and Volkow, 2006; Zacny et al., 2003). Opioids such as oxycodone are mainstays of pain management (Cowan et al., 2003; Minhas and Leri, 2017). but the relatively easy access to these medications and their common diversion has led to a higher incidence of addiction in the exposed population (Cicero and Ellis, 2018, 2017a, 2017b). This increase has deadly consequences; in 2014, oxycodone ranked third behind heroin and cocaine in causing overdose deaths (Warner et al., 2016). As initial use of opioids can result in dependence that leads to concomitant or replacement use of more accessible, less expensive but more lethal opioids, i.e. heroin or fentanyl, it is critical to understand the factors underlying prescription opioid abuse liability (Cicero and Ellis, 2018).

To understand these factors, preclinical researchers would benefit from animal models of prescription opioid abuse liability that recapitulate key aspects of long-term use, such as voluntary and escalating intake that results in tolerance. Prescribed oxycodone is typically taken orally, and animal models of oral opioid intake have been in use for many years (Cappell and LeBlanc, 1971; Enga et al., 2016; Heyne, 1996; Klein, 1995; Nichols et al., 1956; Risner and Khavari, 1973; Shaham et al., 1992; Stolerman and Kumar, 1972; Wikler et al., 1963; Yanaura et al., 1980). Some oral intake paradigms predispose the rodent to opioid drinking via intraperitoneal injection of opioids (Nichols, 1963; Seevers and Deneau, 1963), or force opioid drinking by restricting access to non-drugged water (Cappell and LeBlanc, 1971; Kumar et al., 1968; McMillan et al., 1976; Thompson and Ostlund, 1965). While useful for assessing the impact of opioids on physiology and behavior, such paradigms confound the motivational component that characterizes *voluntary* drug-seeking behavior and opioid reinforcing properties (Meisch, 2001). One classic, true voluntary oral intake model is the “two-bottle choice” paradigm, widely-used in studies of rodent alcohol intake (Belknap et al., 1993; Meisch and Beardsley, 1975; Sinclair, 1976; Taylor et al., 1990; Weiss et al., 1990) but also employed for studies of rodent opiate intake (Hill and Powell, 1976). Whereas some two-bottle studies adulterated the opioid solution with an alternative reinforcer, such as sucrose (Alexander et al., 1981), others found rats would drink an unadulterated opioid solution and even prefer it to water (Heyne, 1996). The use of long-term, free access to an opioid solution in addition to water has provided useful insights, showing, for example, that during periods of controlled drug choice opioid intake can be modified by environmental and individual factors (Heyne, 1996; Pelloux et al., 2006).

Despite this progress, there remain four knowledge gaps in the current literature on long-term, continuous free access to opioids via two-bottle choice. First, morphine and other opioid-agonists have been assessed in two-bottle choice, but oxycodone has not. This is a notable gap in the literature given oxycodone’s distinct pharmacokinetic profile and its non-medical use worldwide. Second, although studies have examined oral opiate intake in the two-bottle choice paradigm, many have used acute, intermittent, or sub-chronic two-bottle access. Long-term continuous access studies can provide additional clinically-relevant insight into the acquisition, maintenance, and potential tolerance to oral opioids. Third, although sex differences in human and rodent oral opioid intake have been long appreciated (Alexander et al., 1978; Craft, 2008; Fillingim and Gear, 2004; Graziani and Nisticò, 2016a; Hadaway et al., 1979; Klein, 1995; reviewed in Serdarevic et al., 2017), female rodents have not been assessed in a paradigm that affords long-term, continuous access to oral oxycodone. Fourth, although pre-exposure endophenotypes can predict later opioid use (such as high vs. low locomotor activity in a novel environment), such pre-screening has not been performed for oxycodone in a long-term voluntary choice paradigm.

The development of a long-term voluntary choice opioid oral intake model would complement other rodent models of dependence and addiction, such as IV self-administration, thereby facilitating intervention testing. Here we employ a classic 2-bottle choice paradigm to provide female and male rats with long-term, free, and continuous access to an oxycodone solution and water. Adult experimental female and male Long Evans hooded rats were given free access (24 h/d, 7 d/wk) to a bottle containing oxycodone and a bottle of plain water in their home cage (Experimental rats), whereas Control rats had access to two bottles of water. Most Experimental rats spontaneously and exclusively drank oxycodone, escalating their intake over time, and female rats drank more compared to male rats. Motivation to self-administer oxycodone (via citric acid suppression test) revealed that both Experimental and Control rats suppressed their intake of citric acid-adulterated oxycodone or water, respectively. However, Experimental rat oxycodone intake immediately returned to pre-citric acid intake levels, whereas Control rat water intake did not. Oxycodone dependence via naloxone-precipitated withdrawal was evident in Experimental but not Control rats. Overall, this model recapitulates many features of human oxycodone abuse in female and male rats and will be a useful addition to the field for probing the neural and physiological mechanisms that mediate long-term voluntary drug use and exploration of potential treatment options.

## 2. MATERIAL AND METHODS

### 2.1. Animals and husbandry

Female (n=12) and male (n=12) Long-Evans Hooded rats were purchased from Envigo (formerly Harlan) at the age of 60-65 days. They were individually housed in a vivarium with constant temperature 20-23°C and humidity 30-70% and *ad libitum* food and water available via lixts until after behavioral prescreening (see section **2.2**) was complete. The lights were turned off at 6PM and turned on at 6AM. Rats were handled 3 times per week for 2-3 minutes to familiarize them with different experimenters. Rat weights were recorded once before behavioral testing and once a week throughout the experiment until terminal procedures were performed. The cage was removed and cleaned once a week by experimenters (not by husbandry staff), with an occasional spot change for cages deemed unsanitary. Corn cob bedding was supplemented with paper shreddings and a plastic tunnel.

### 2.2. Baseline behavioral assessment

To prescreen rats, after ∼2 weeks of individual housing rats were tested in the open field and for marble burying behavior **(Fig. 1)**. The data from these tests were used to 1) assure that the random assignment to subsequent Experimental and Control groups was unbiased for the traits measured; and 2) to determine if the behaviors from these tests could be used to predict later outcome measures once oxycodone was available. Open field was also repeated during Week 31 as postscreening evaluation.

**Figure 1.**
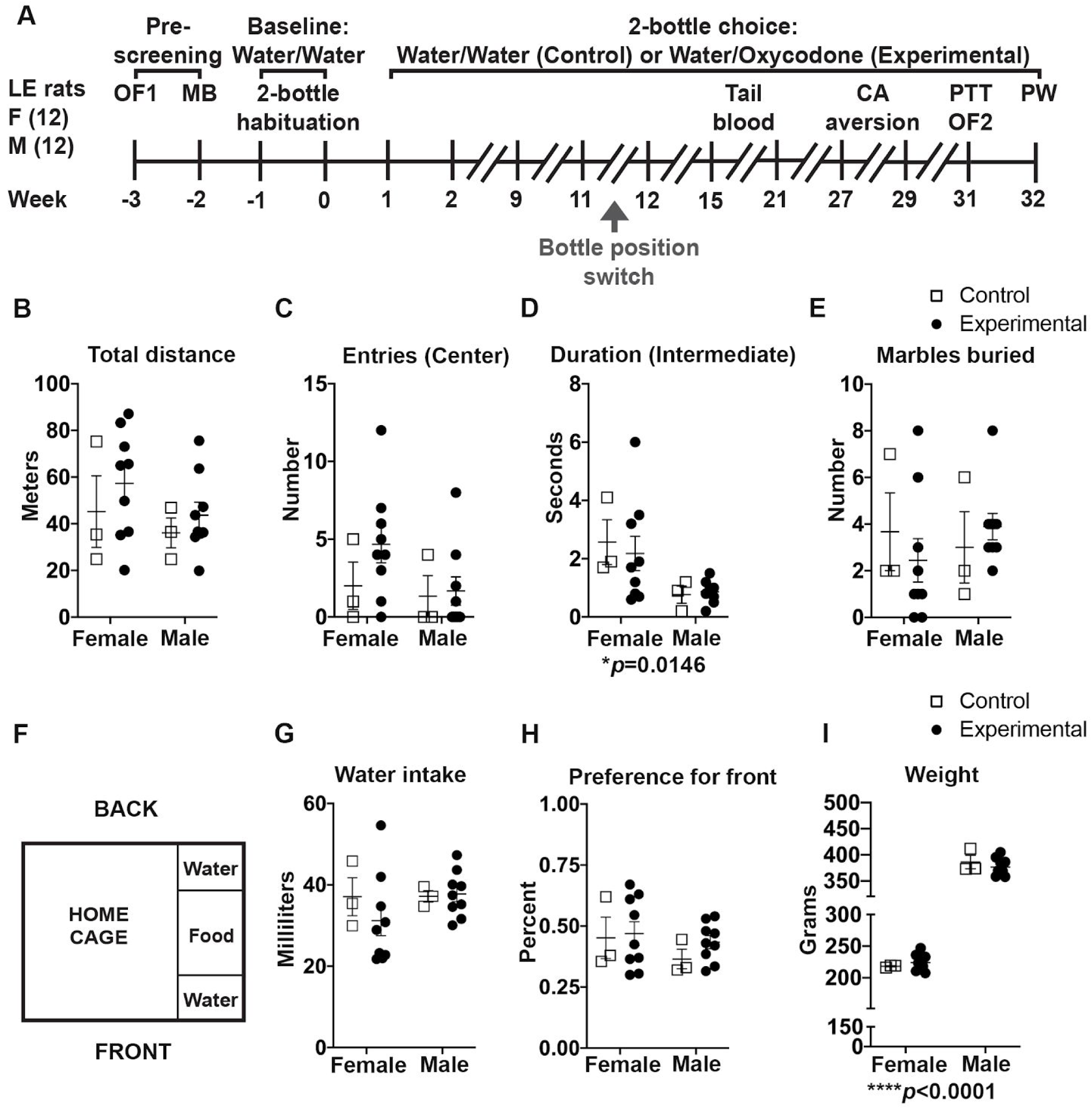
Pre-screening behavioral assessment (before group assignment) revealed no overt group differences. **(A)** Experimental time course. Long Evans (**LE**) hooded rats, 12 females (**F**) and 12 males (**M**), were **pre-screened** (Weeks −3 and −2) on the open field **(OF1)** and marble burying **(MB)** tests. After pre-screening, all rats entered the **Baseline** (**Water/Water**) period where they were habituated for two weeks (Weeks −1 and 0) to drink from two water bottles, one each in front of or behind the food hopper positioned on the right side of the cage (see panel **1F**). Starting at Week 1, **Control** rats (3 female and 3 male rats, randomly chosen) continued to receive two bottles containing water (**Water/Water**); **Experimental** rats (9 female and 9 male rats, also randomly chosen) received an oxycodone-containing bottle in the front position and a water-containing bottle in the back position (**Water/Oxycodone**). At the end of Week 11/start of Week 12, the front bottle (containing water or oxycodone) was switched to the back position for both Control and Experimental rats (**Bottle Position Switch**). About 8 hours into the 12-hour dark cycle, during Weeks 15 and 21, tail vein blood was extracted from all rats. From Weeks 27 to 29, all rats were intermittently exposed to increasing concentrations of citric acid (**CA**) to test for aversion. Week 31, all rats were tested for nociception in the plantar thermal test (**PTT**) and open field again (**OF2**) subsequently for signs of naloxone precipitated withdrawal (**PW**). **(B)** The analysis of total distance traveled over the 30 min in the OF1 via a two-way ANOVA (Sex x Treatment) revealed no interactions, F(1,20)=0.05, *p*=0.81, no main effect of Sex, F(1,20)=1.41, *p*=0.24, and no main effect of Treatment, F(1,20)=1.06, *p*=0.31. Note that in this panel **B** and panels **C-E** and **G-I**, Experimental and Control refers to the subsequent assignment of naive rats to these two groups. **(C)** Analysis of number of OF1 entries into the Center area using a two-way ANOVA (Sex x Treatment) revealed no interaction, F(1,20)=0.66, *p*=0.42, no main effect of Sex, F(1,20)=1.63, *p*=0.21, and no main effect of Treatment, F(1,20)=1.09, *p*=0.30. **(D)** Analysis of duration of time spent in the OF1 Intermediate area by a two-way ANOVA (Sex x Treatment) revealed no interactions F(1,20)=0.17, *p*=0.67, a significant effect of Sex F(1,20)=7.14, **\**p*=0.015**, and no main effect of Treatment F(1,20)=0.06, *p*=0.80. **(E)** Marble burying test analysis of the number of marbles buried using a two-way ANOVA (Sex x Treatment) revealed no interaction F(1,20)=0.86, *p*=0.36, no main effect of Sex F(1,20)=0.11, *p*=0.73, and no main effect of Treatment F(1,20)=0.02, *p*=0.88. **(F)** Schematic of the home cage during the two-bottle habituation (Baseline) period, with water (W) bottles placed in the front and the back of the food hopper on the right side of the cage. Rats had access to both bottles 24 hours/day and 7 days/week. Not drawn to scale. **(G)** Analysis of total water intake during the Baseline period via two-way ANOVA (Treatment x Sex) revealed no interaction, F(1,20)=0.67, *p*=0.42, no main effect of Sex, F(1,20)=0.71, *p*=0.40, and no main effect of Treatment, F(1,20)=0.45, *p*=0.50. **(H)** Analysis of position preference (ratio of intake from the front bottle over the total intake) during the Baseline period via two-way ANOVA (Treatment x Sex) revealed no interaction, F(1,20)=0.22, *p*=0.64, no main effect of Sex, F(1,20)=1.22, *p*=0.28, and no main effect of Treatment, F(1,20)=0.61, *p*=0.44. **(I)** Analysis of body weight during the Baseline period via two-way ANOVA (Treatment x Sex) revealed no interaction, F(1,20)=1.16, *p*=0.29, a significant main effect of Sex, F(1,20)=503.2, **\*\*\*\**p*<0.0001**, and no effect of Treatment, F(1,20)=0.06, *p*=0.79.

The open field was performed as previously described (Bouwknecht et al., 2007; Kulesskaya and Voikar, 2014) with minor modifications. The open field was conducted in a square-shaped arena (76 × 76 × 30 cm) with white plastic floor and white walls. The following behavioral parameters were scored using ANY-maze software: total distance traveled, average speed, maximum speed in Center and Intermediate zones of the open field, longest time period spent in the Center and Intermediate zones, number of entries in Center and Intermediate zones, duration of time in the Center and Intermediate zones, average duration in the Center and Intermediate zones, latency to enter the Center and Intermediate zones, and number of fecal boluses deposited during test. One day later, rats were then assessed for marble burying, as previously described (Deacon, 2006; Ku et al., 2016; K. ‘u Njung’e and Handley, 1991; Schneider and Popik, 2007; Thomas et al., 2009a) with minor modifications. Briefly, rats were habituated for 10min in a cage filled with approximately 5cm corn cob bedding. The rat was removed and 18 marbles (1cm diameter each) were placed in the cage in a 3×6 pattern. The rat was then reintroduced in the cage for 20min period of testing. Three blinded observers scored the number of marbles buried, and a consensus was reached after visual inspection of each cage with consultation from the offline video score as-needed.

### 2.3. Oxycodone self-administration

After behavioral assessment **(Fig. 1)**, lixits were removed from all cages and rats were given water via a bottle (Hydropac® or Zyfone bottle, as described below) for the duration of the experiment. Intake of liquid was measured at 7AM and 4PM each day until Week 11, and at only 7AM from Week 11 until the end of the experiment. Liquid intake was calculated as the change in weight (g) of the bottle from the previous measurement. A ratio was calculated as the amount of liquid drunk (g) from one of the two liquid bottles over the total intake from the two bottles. During Baseline (Weeks −1 and 0, **Fig. 1)**, rats were provided two water Hydropacs (in front and back position) in a hopper on the right side of the cage with food placed in between. Starting on Week 1, Control (“Water/Water”) rats continued to receive water from both bottles, whereas Experimental (“Water/Oxycodone”) rats received water in one bottle and oxycodone dissolved in water in the other. The starting oxycodone concentration in each bottle was determined for each rat based on their baseline average water intake and their baseline body weight. This allowed the animals to titrate their own intake, and resulted in a delivered concentration ranging from 0.06 to 0.12 mg/mL, with a goal of individual rat intake of ∼10 mg/kg/day (Schrott et al., 2008) if they exclusively drank from the oxycodone bottle. From Weeks 3 to 11, the oxycodone bottle was placed in the front hopper position, and from Weeks 12 to 32 it was switched to back hopper position (see “Bottle position switch”, **Fig. 1A**). At Week 8, Hydropacs were replaced by water bottles (ZyfoneTM One CageTM 16 oz) with a twist cap (ZyfoneTM high-temperature plastic One CapTM) and a 1.5” Sipper Tube With Stainless Ball (Part#-ST3.0SB, Alternative Design Mfg & Supply Inc.) due to extensive chewing of the Hydropac plastic valve and spillage by some rats.

### 2.4. Home cage behaviors

Home cage behaviors were observed twice daily, at 7AM and 4PM. A single rat was tracked by one observer during 1-min observation periods for the presence or absence of specific behaviors **(Table 1)** indicative of normal activity (i.e. grooming, walking) as well as spontaneous withdrawal-like behavior (i.e. wet dog shakes) as adapted from previous studies (Cicero et al., 2002a; Gellert and Holtzman, 1978a; Jones and Barr, 1995; Koob et al., 1992; Rasmussen et al., 1990).

**Table 1.** Spontaneous and precipitated withdrawal behaviors.

| **Spontaneous and precipitated withdrawal behaviors** |  |  |
| --- | --- | --- |
| **Behavior** | **Definition** | **How recorded?** |
| Head movement | Lateral and rotary motion of the head | 0≥1 with +1 if occurs >1X |
| High walking | Elevated torso or spine curved upwards, can present as abnormal posture in rats. | 0≥1 with +1 if occurs >1X |
| Jumping | Sudden leaping such that all four paws are off the bottom of the chamber | 0≥1 with +1 if occurs >1X |
| Ptosis | One or both eyelids closed or semi-closed | 0 = absent; 1 = mild; 2 = moderate; 3 = marked |
| Wet dog shakes | Rapid shaking of the whole body | 0≥1 with +1 if occurs >1X |
| Burrow | Sliding the head under the corn cob cage bedding | 0≥1 with +1 if occurs >1X |
| Diarrhea | Extremely loose and watery stool | 0 = absent; 1 = mild; 2 = moderate; 3 = marked |
| Yawning | Wide opening and closing (stretching) of the mouth. Can be seen with and without body stretching (separate behavior). | 0≥1 with +1 if occurs >1X |
| Stretching | Extension or dorsal flexion of the trunk causing a visual lengthening of the body | 0≥1 with +1 if occurs >1X |
| Teeth chatter | Movement of the jaw with constant chattering of the teeth, or distinct repetitive movement of cheek/facial muscles. | 0≥1 with +1 if occurs >1X and each occurrence separated by 3 seconds |
| **Only precipitated withdrawal behaviors** |  |  |
| **Behavior** | **Definition** | **How recorded?** |
| Writhing | Abdominal stretching | 0≥1 with +1 if occurs >1X |
| Sweeping tail | Heavily carrying the tail around | 0≥1 with +1 if occurs >1X |
| Chewing | Without matter in the mouth | 0≥1 with +1 if occurs >1X |
| Ejaculation | Releasing semen | 0 = absent; 1 = mild; 2 = moderate; 3 = marked |
| Piloerection | Bristling of hair | 0 = absent; 1 = mild; 2 = moderate; 3 = marked |
| Straub tail | Curved upwards tail | 0≥1 with +1 if occurs >1X |

### 2.5. Serum assessment of oxycodone levels

Blood was taken via the tail vein under isoflurane anesthesia. Blood was collected in sterile Eppendorf tubes with 10µl of heparin and centrifuged at 4K RPM for 10min and serum separated and stored at −20°C. Serum was sent to the Bioanalytical Core Center for Clinical Pharmacology at the Children’s Hospital of Philadelphia Research Institute. Assay development consisted of oxycodone separation through ultra-performance liquid chromatography (UPLC) and detection by tandem mass spectrometry (MS-MS) using a triple quadrupole mass spectrometer (API4000).

### 2.6. Gnawing home cage behavior

At the start of Week 12, each rat cage received 1 nylabone and 1 wood gnawing block (Bio-Serv K3512 medium size). The weight of both the nylabone and block were recorded daily at the same time of day. The change in weight of each was calculated to derive a gnawing index, defined as the total amount in grams of nylabone and wood block chewed over time.

### 2.7. Citric acid aversion test

Between Week 27 and 29 all rats received ascending concentrations of citric acid (CA, Sigma Aldrich, cat# PHR1071) at 1, 3, and 5mM in the back bottle. 5mM CA was repeated once (5 and 5’). In between each CA exposure, rats received their original bottle (water or oxycodone) for 2 days to allow for a return to baseline. Liquid intake, body weight, preference, and deviation from baseline were scored daily.

### 2.8. Plantar thermal test

On Week 31 all rats were tested for analgesia in the plantar thermal test using a Hargreaves apparatus (IITC Inc. Life Science, Model 390G, Series 8, Heated Base) as previously described (Barr et al., 2016). Rats were habituated for 10min in a custom acrylic enclosures with high walls placed on the tempered (30°C) glass surface of the apparatus. The test was performed on each hindpaw plantar surface by a focused, radiant heat light source creating a 4X6mm focal spot set to a measured temperature of 60°C which in preliminary work with other animals produced a baseline withdrawal latency of ∼10-14s. We used a cut-off of 20s if the animal did not respond, avoiding tissue damage. Each hindpaw was measured 3 times with approximately 1-2min between tests.

### 2.9. Precipitated withdrawal

On Week 32 all rats were tested for signs of withdrawal. Rats were habituated to a clean cage containing corn cob bedding for 10min and behavior recorded and scored for total distance traveled using ANY-maze software and 3 side cameras to capture closely signs of withdrawal. After habituation, each rat was injected with naloxone (1mg/kg, i.p. Sigma-Aldrich cat# PHR1802) and scored for 15min for signs of withdrawal. Withdrawal symptoms were scored as previously described and summarized in **Table 1** (Bläsig et al., 1973; Cicero et al., 2002b; Gellert and Holtzman, 1978b; Harvey-Lewis et al., 2015; Koob et al., 1992; Rasmussen et al., 1990). Weight was measured before habituation and after 15min of precipitated withdrawal. One weight data point for one animal was lost.

### 2.10. Statistical analysis

Data analysis was performed using GraphPad Prism® (version 8; Macintosh) or PAST3 (version 3.2). A two-tailed unpaired t-test was used to test the difference of the means between two groups. A 2-way repeated measures (RM) ANOVA or a mixed-effects model (REML) tested for main effects of Treatment or Sex and their interaction. If an interaction was significant after analysis of variance, a post-hoc test for multiple comparisons correction was performed. Dunnett’s post-hoc test was performed when comparisons were made to a single control group; Sidak’s or Tukey’s tests were performed for multiple comparisons between different groups. Data are presented as individual data points and as mean ± SEM. In all analyses, the level of significance was *p*<0.05. Exact statistical results and *p* values for the statistical tests are provided in figure panels and legends.

A multivariate sample-based factor analysis (PAST3; Q-Mode CAB-FAC (Imbrie and Kipp, n.d.)) was conducted separately on the pre- and post-screening open field and marble burying behaviors for females and males, obtaining a single score for each rat, by multiplying the pre-screening open field behavior measures by the factor score of that behavior. That summed score was then correlated with the dependent measures during the intake phase (precipitated withdrawal behaviors, spontaneous withdrawal behaviors, dose, intake, preference, citric acid, blood, gnawing) for the two independent data sets to obtain the best-fit value of the slope (PAST3; r^2^).

## 3. RESULTS

### 3.1. Pre-screening for a predictive endophenotype: assessment prior to experimental assignment revealed no overt group differences before group assignment

#### 3.1.1. Pre-screening behavioral assessment

Before any manipulations (Week −3 and Week −2; **Fig. 1A**), rats were pre-screened for open field (OF1) behavior, as an indicator of both anxiety and locomotor activity (Colman, 2001; Markel’ et al., 1989) and for marble burying, as a repetitive behavior (Thomas et al., 2009b) often associated with an anxiety-like phenotype (K. Njung’e and Handley, 1991). For OF1 the total distance traveled **(Fig. 1B)** and the number of entries into the Center zone **(Fig. 1C)** were similar within groups irrespective of Sex or the to-be-assigned Treatments (Control or Experimental). There was a significant main effect of Sex of duration in the Intermediate zone **(Fig. 1D)**, with females spending ∼3x more time in the Intermediate zone than males. We found no difference in the number of marbles buried driven by Sex or to-be-assigned Treatment groups **(Fig. 1E)**. Thereby at pre-screening, the behavior parameters were similar, with the exception that females showed higher levels of novelty-seeking (longer time spent in the Intermediate zone in the OF1 relative to males).

#### 3.1.2. Baseline Water/Water intake

At Baseline (average of Week −1 and Week 0, **Fig. 1A**) two bottles of water were provided to both Control and Experimental rats in their home cage to allow for habituation **(Fig. 1F)**. There was no effect of Treatment or Sex in the average amount of liquid ingested over the first two weeks of habituation **(Fig. 1G)**. There was no effect of Treatment or Sex in the preference for the front or the back bottle **(Fig. 1H)**. Within Sex, rats had similar body weights **(Fig. 1I)**, with females weighing, as expected, less than males. Thus, the random assignment to either Control or Experimental group resulted in unbiased inclusion in the two groups.

### 3.2. Experimental rats increased their oxycodone preference and drank more oxycodone over time, with females ingesting more than males with resultant higher oxycodone blood levels

#### 3.2.1. Two bottle-choice period

During the 2 bottle-choice period, female Experimental rats increased their total liquid intake from Week 19 compared to Week 1, whereas male Experimental rats did not at any week **(Fig. 2A)**. Both females and males increased their preference for oxycodone by Week 9 relative to Week 1. When the bottle position was switched, both females and males maintained their preference for oxycodone, but male rats then increased their preference for oxycodone by Week 17 compared to the week before the bottle switch (Week 11; **Fig. 2B**). In contrast, controls had a stable intake of water **(Fig. 2C)** and preference for the bottle position (slightly preferring the back bottle; **Fig. 2D**).

**Figure 2.**
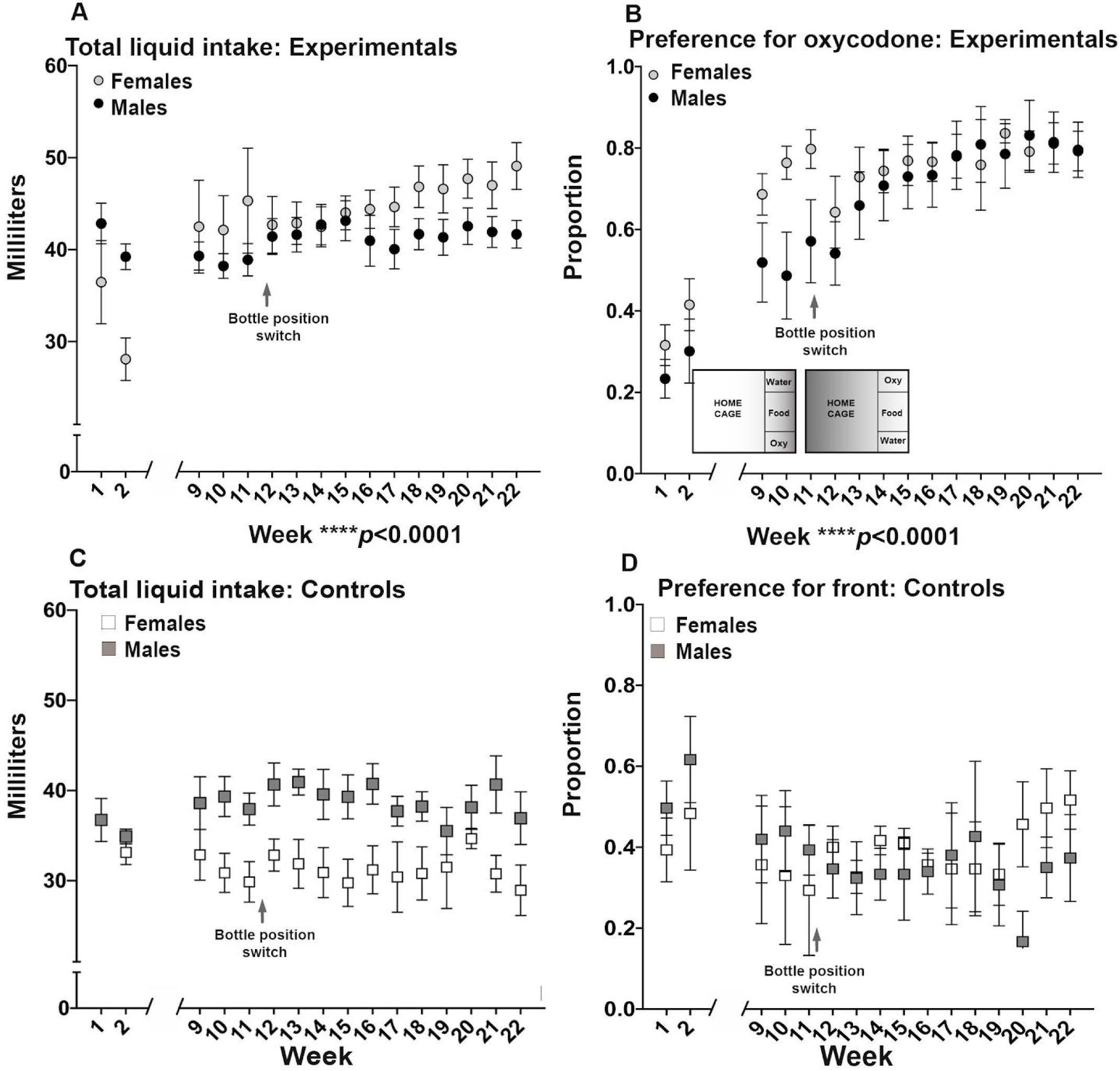
Experimental rats increase liquid intake and oxycodone preference over time in a sex-dependent manner. **(A)** Total liquid intake in Experimental rats (receiving Water/Oxycodone beginning Week 3) analyzed via a restricted mixed-effects model (Sex x Time) revealed a significant interaction, F(15,222)=2.942, **\*\*\**p*=0.0003**, no main effect of Sex, F(1,15)=0.7735, *p*=0.3930, and a significant effect of Time, F(15, 222)=4.153, **\*\*\*\**p*<0.0001**. Post-hoc analyses within each Sex showed that compared to Week 1, males did not change their liquid intake over time, whereas females did (Week 1 vs. Week 18 ***p*=0.0035**, Week 1 vs. Week 19 ***p*=0.0049**, Week 1 vs. Week 20 ***p*=0.0010**, Week 1 vs. Week 21 ***p*=0.0028**, Week 22 ***p*=0.0001**). Post-hoc analysis of total intake between females and males showed no sex difference for any given week. **(B)** Oxycodone preference of Experimental rats analyzed via a restricted mixed-effects model (Time x Sex) revealed a significant interaction F(15,222)=1.990, **\**p*=0.00170**, no main effect of Sex F(1,15)=0.6337, *p*=0.4384, and a significant effect of Time F(15, 222)=22.78, **\*\*\*\**p*<0.0001**. Post-hoc comparison within Sex showed that both females and males increased their preference by Week 9 relative to Week 1 (females: Week 1 vs. Weeks 9-22 ***p*<0.0001**; males: Week 1 vs. 9-22 ***p*<0.0001**). When the bottle position was switched, post-hoc analyses revealed both females and males maintained their preference for the oxycodone bottle (Week 11 vs. all the subsequent post-bottle switch weeks did not differ). Male rats then escalated their preference for oxycodone by Week 17 relative to Week 11 (Week 11 vs. 17 ***p*=0.0112**, vs. 18 ***p*=0.0026**, vs. 19 ***p*=0.0095**, vs. 20 ***p*=0.0007**, vs. 21 ***p*=0.0019**, vs. 22 ***p*=0.0056**), whereas females did not. The inset shows the position of water and oxycodone (**Oxy**) bottles in Experimental rat cages until Week 11 and after bottle position switch at Week 12. **(C)** Total liquid intake in Control rats (receiving Water/Water throughout time) analyzed via two-way RM ANOVA (Sex x Time) revealed a significant interaction, F(15,60)=2.492, **\*\**p*=0.0064**, no main effect of Sex, F(1, 4)=5.156, *p*=0.0857, and no effect of Time, F(3.085,12.34)=1.656, *p*=0.2271. Post-hoc analyses within each Sex showed that compared to Week 1, neither females nor males t differed over time in their liquid intake. Post-hoc analysis of total intake between females and males showed no difference at any given week. **(D)** Preference for front bottle in Control rats (receiving Water/Water throughout time) analyzed via two-way RM ANOVA (Sex x Time) revealed no interaction, F(18,72)=0.8038, *p*=0.6895, no main effect of Sex, F(1, 4)=0.002583, *p*=0.9619, and no significant effect of Time, F(18,72)=0.9619, *p*=0.9619.

#### 3.2.2. Body weight over time

The body weights of female Control and female Experimental rats did not differ from each other at any time point. Male Experimental rats weighed significantly less than Control male rats from Week 13 onwards **(Fig. 3A)**. We calculated both the absolute intake (mg) and intake by body weight (mg/kg) for each animal. Female and male rats self-administered both a higher dose of oxycodone over time by the absolute amount and by mg/kg, with females self-administering a higher dose by Week 9, whereas males self-administered a higher dose by Week 14 **(Figs. 3B, 3C)**. Females self-administered higher doses per body weight than did males on Weeks 9 to 12, 17, 18 and 20 to 22 **(Fig. 3C)**. The greater oxycodone intake in females was reflected in higher oxycodone blood levels compared to males (**Fig. 3D**, left panel). When the dose of oxycodone was correlated with blood levels, the relationship was significant and positive for males, but not significant for females (**Fig. 3D**, right panel).

**Figure 3.**
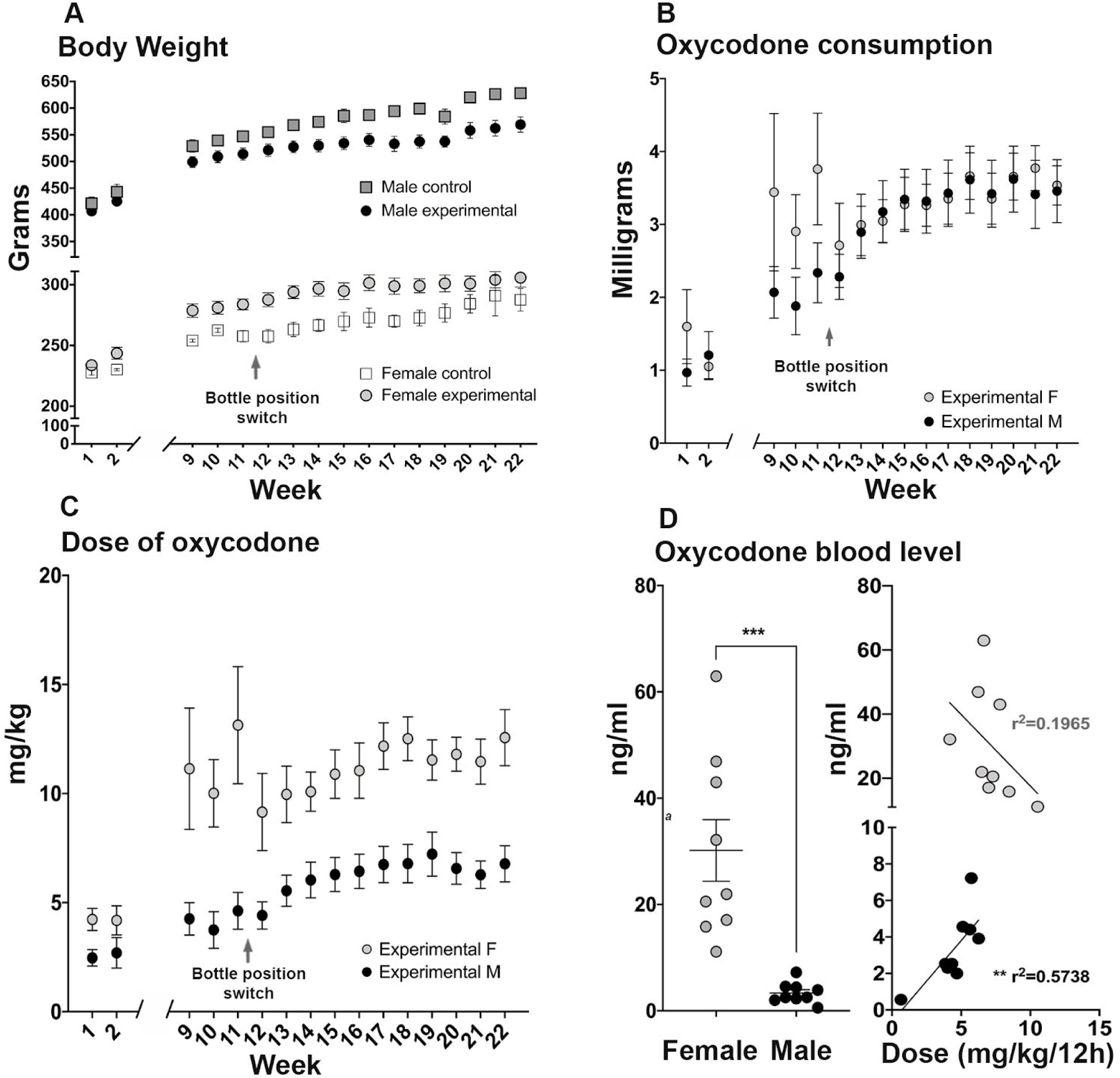
Experimental rats intake increasing amounts of oxycodone with resultant high blood levels of the drug, especially for females. **(A)** Body weights over time in Control and Experimental rats analyzed via three-way ANOVA (Time x Treatment x Sex) revealed a significant effect of Time F(3.615, 68.68)=488.1, **\*\*\*\**p*<0.0001**, a significant effect of Sex, F(1,19)=577.0, **\*\*\*\**p*<0.0001**, and no effect of Treatment, F(1,19)=0.2491, *p*=0.6235. The three-way interaction (Sex x Treatment x Time) was significant, F(23, 437)=7.825, p<.0001 and posthoc tests showed that female Control rats and Experimental rats did not differ at any time point. Male Experimental rats weighed significantly less than Male Control rats from Week 13 onwards. **(B)** Analysis of self-administered oxycodone intake (mg) via a mixed-effects model (Time x Sex) revealed no significant interaction F(15,223)=1.641, *p*=0.0648, a significant main effect of Time F(15,223)=12.84, **\*\*\*\**p*<0.0001**, and no main effect of Sex F(1, 15)=0.4683, *p*=0.5042. Posthoc analyses of the main effect of time showed that all weeks differed from weeks 1 and 2 but that no weeks from 9 to 22 differed from week 11. Thus for females and males intake was steady across time. **(C)** Analysis of self-administered dose via a mixed-effects model (Time x Sex) revealed no significant interaction F(15,222)=1.711, *p*=0.0502, a significant main effect of Sex F(1,15)=18.72, \*\*\****p*=0.0006**, and a significant effect of Time F(15, 222)=10.04, **\*\*\*\**p*<0.0001**. Post-hoc analysis within same sex group showed that female rats self-administered oxycodone at a higher dose from Week 9 on compared to Week 1 (Week 1 vs 10 ***p*=0.0002**, Week 1 vs 12 ***p*=0.0030**, Week 1 vs 13 ***p*=0.0002**, Week 1 vs 14 ***p*=0.0001**, and all the other individual weeks compared to Week 1, ***p*<0.0001**). Males self-administered oxycodone at a higher dose from Week 14 on compared to Week 1 (Week 1 vs 14 ***p*=0.0140**, week 1 vs 15 ***p*=0.0065**, week 1 vs 16 ***p*=0.0041**, week 1 vs 17 ***p*=0.0015**, week 1 vs 18 ***p*=0.0012**, week 1 vs 19 ***p*=0.0003**, week 1 vs 20 ***p*=0.0026**, week 1 vs 21 ***p*=0.0067**, week 1 vs 22 ***p*=0.0013**). Post-hoc analysis of oxycodone dose of female compared to male rats at each time point showed a significant difference at Weeks 9 to 12 compared to week 1 (Week 9 ***p*=0.0004**, Week 10 ***p*=0.0017**, Week 11 ***p*<0.0001**, Week 12 ***p*=0.0500**), Weeks 17 and 18 (Week 17 ***p*=0.0120**, Week 18 ***p*=0.0062**), and Weeks 20 to 22 (Week 20 ***p*=0.0184**, Week 21 ***p*=0.0204**, Week 22 ***p*=0.0054**). **(D) Left panel:** Analysis via two-tailed unpaired t-test of oxycodone blood levels between female and male rats averaged over the two blood draws at Weeks 15 and 21 revealed a significant difference between sexes (**\*\*\**p*=0.0003**). **Right panel:** Correlation analysis of the blood levels with the intake 12 hours before blood collection revealed a significant positive correlation in males with the slope of the linear regression of 0.8874, r^2^=0.5738, and ***p*=0.0181**, whereas for females the slope of the linear regression was −4.458, r^2^=0.1965, and *p*=0.2320.

### 3.2. Experimental animals show higher levels of dependency in precipitated withdrawal and increased gnawing behavior but no changes in nociception

#### 3.2.1. Withdrawal in the home cage

Daily assessment of home cage spontaneous withdrawal symptoms (burrow, teeth chatter, stretch, and jump) at 7AM showed no significant difference between groups when analyzed as individual behaviors or when the total sum of these behaviors was collapsed over 22 weeks **(Fig. 4A)**. Overall, Experimental rats chewed more nylabone and wooden block in the home cage between Weeks 12 and 22 of oxycodone exposure than did Control rats; moreover, Experimental females gnawed more than Control females, Control males, and Experimental males. Experimental males did not differ from Control males **(Fig. 4B)**.

**Figure 4.**
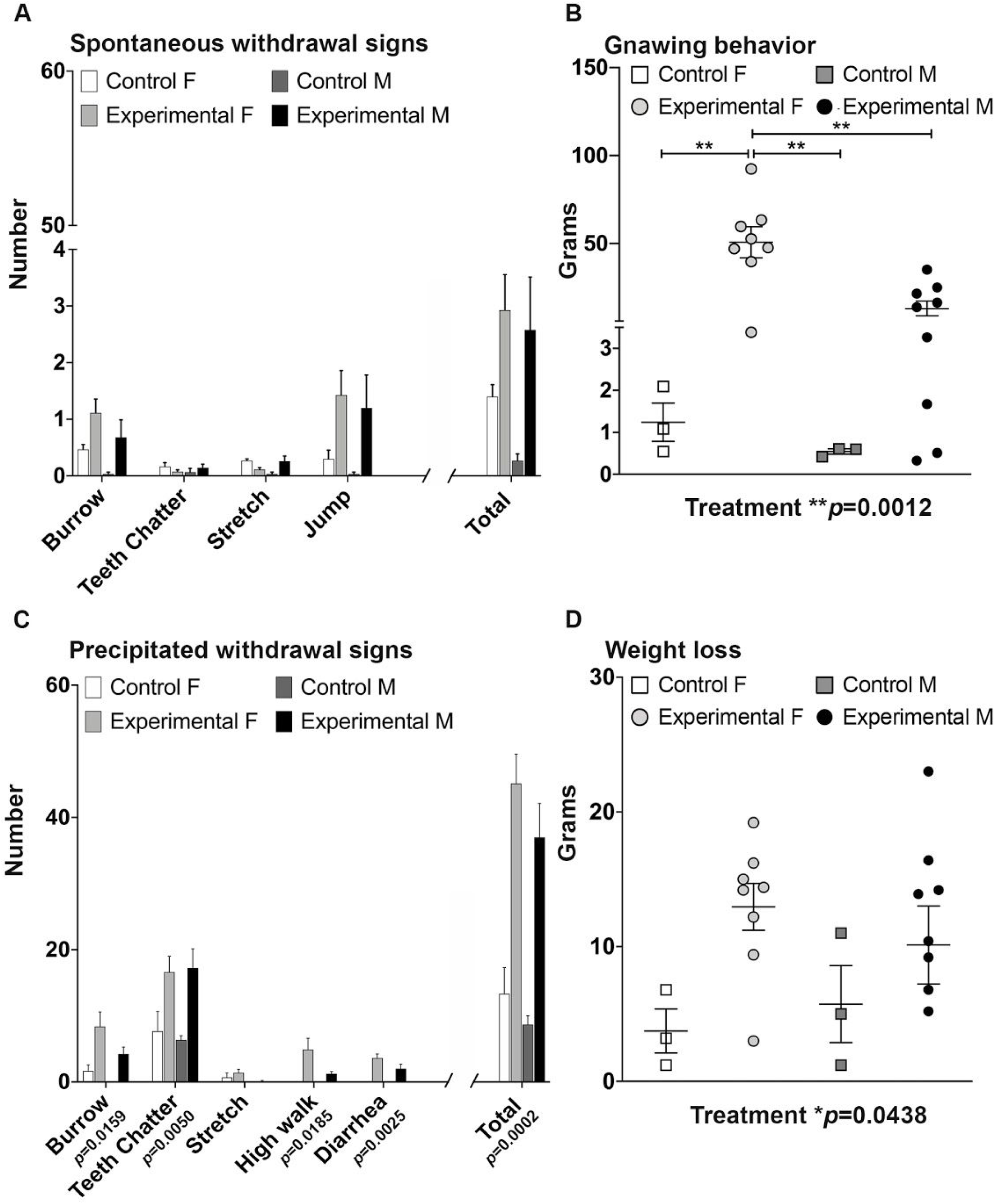
Withdrawal signs gnawing behavior and are increased in experimental rats - particularly in females - relative to control rats. **(A)** Individual spontaneous withdrawal sign analysis via two-way ANOVA (Sex x Treatment) revealed no interaction and no effect of Treatment or Sex for frequently occurring behaviors: **Burrow** (interaction, F(1,19)=3.935^e-006^, *p*=0.9984; Sex, F(1,19)=1.537, *p*=0.2302; Treatment, F(1,19)=3.396, *p*=0.0810), **Teeth chatter** (interaction, F(1,19)=1.547, *p*=0.2287; Sex, F(1,19)=0.05031, *p*=0.8249; Treatment, F(1,19)=0.01040, *p*=0.9199), **Stretching** (interaction, F(1,19)=4.006, *p*=0.0598; Sex, F(1,19)=0.2305, *p*=0.6366; Treatment, F(1,19)=0.1310, *p*=0.7214), **Jumping** (interaction, F(1,19)=0.001045, *p*=0.9745; Sex, F(1,19)=0.1455, *p*=0.7071; Treatment, F(1,19)=3.161, *p*=0.0914). Two-way ANOVA of the **Total** spontaneous withdrawal signs also showed no interaction F(1,19)=0.1533, *p*=0.6997 or effect of Sex F(1,19)=0.5439, *p*=0.4698 or Treatment F(1,19)=3.651, *p*=0.0712. **(B)** The sum of total amount in grams of chewed nylabone and wooden block averaged over 11 weeks (Weeks 12 to 22). Two-way ANOVA (Treatment x Sex) revealed an interaction F(1,19)=5.106, **\**p*=0.0358**, an effect of Sex F(1,19)=5.499, **\**p*=0.0300**, and an effect of Treatment F(1,19)=14.39, **\*\**p*=0.0012**. Post-hoc analysis within and between Sex revealed experimental females gnawed more than control females **\*\**p*=0.0026**, control males **\*\**p*=0.0023** and experimental males **\*\**p*=0.0015**. **(C)** Individual precipitated withdrawal signs analysis via two-way ANOVA (Sex x Treatment) revealed no interaction and no effect of Sex but a significant effect of Treatment for the following behaviors: **Burrow** (interaction, F(1,19)=0.2538, *p*=0.6202; Sex, F(1,19)=1.821, *p*=0.1931; Treatment, F(1,19)=7.000, ***p*=0.0159**); **Teeth chatter** (interaction, F(1,19)=0.0002038, *p*=0.9645; Sex, F(1,19)=0.1217, *p*=0.7311; Treatment, F(1,19)=10.06, ***p*=0.0050**); **High walking** (interaction, F(1,19)=1.797, *p*=0.1959; Sex, F(1,19)=1.797, *p*=0.1959; Treatment, F(1,19)=6.634, ***p*=0.0185**); **Diarrhea** (interaction, F(1,19)=1.121, *p*=0.3030; Sex, F(1,19)=1.121, *p*=0.3030; Treatment, F(1,19)=12.10, ***p*=0.0025**). However, there was no significant interaction interaction and no effect of Sex or Treatment for **Stretching** (interaction, F(1,19)=0.4660, *p*=0.5031; Sex, F(1,19)=3.775, *p*=0.0670; Treatment, F(1,19)=0.7970, *p*=0.3832). Analysis of the **Total** withdrawal signs via Two-way ANOVA showed no interaction F(1,19)=0.1784, *p*=0.6805, no effect of Sex F(1,19)=1.248, *p*=0.2779 but a significant effect of Treatment F(1,19)=21.73, ***p=*0.0002**. **(D)** Weight loss after precipitated withdrawal analyzed via two-way ANOVA (Sex x Treatment) revealed no interaction F(1,19)=0.5872, *p*=0.4529, no Sex effect F(1,19)=0.01726, *p*=0.8969, but a Treatment effect F(1,19)=4.664, **\**p*=0.0438**.

#### 3.2.2. Precipitated withdrawal

In contrast to home cage spontaneous withdrawal behaviors, Experimental rats presented more precipitated withdrawal symptoms than Control rats at Week 31 for the following discrete behaviors analyzed separately: burrow, teeth chatter, high walk, and diarrhea. When the total precipitated withdrawal score was analyzed, Experimental females and males presented more withdrawal signs than their respective controls **(Fig. 4C)**. After precipitated withdrawal, Experimental rats lost more weight than did Controls Experimental females lost four-fold more weight than Control females and Experimental males lost two-fold more than Control males **(Fig. 4D)**.

#### 3.2.3. Plantar thermal test

When measuring paw withdrawal latency in the plantar thermal test, there was no effect of Treatment or Sex and no Treatment by Sex interaction. The means and SEM (in seconds) for the four groups were female Control: 13.93 ± 2.29; female Experimental: 12.53 ± 1.9; male Experimental: 11.14 ± 1.14; male Control: 14.74 ± 1.75. Moreover, the correlations between oxycodone intake the day before the plantar test were non-significant for females, males or the two sexes combined (r’s = 0.019; 0.356; 0.267 respectively).

### 3.3. Increasing concentration of CA suppressed intake in both Control and Experimental rats, but Experimental rats returned to initial baseline preference whereas Control rats did not

When CA was added to the back bottle **(Fig. 5A)**, all rats suppressed their intake from that bottle at higher CA concentrations compared to their intake the day before the first citric acid test (BL0; **Fig. 5B**). Experimental males suppressed at 3mM CA whereas the other groups suppressed at 5mM **(Fig. 5B)**. When CA was removed during the interspersed baselines, Experimental rats returned to the initial baseline preference, whereas Control rats maintained their aversion **(Fig. 5C)**.

**Figure 5.**
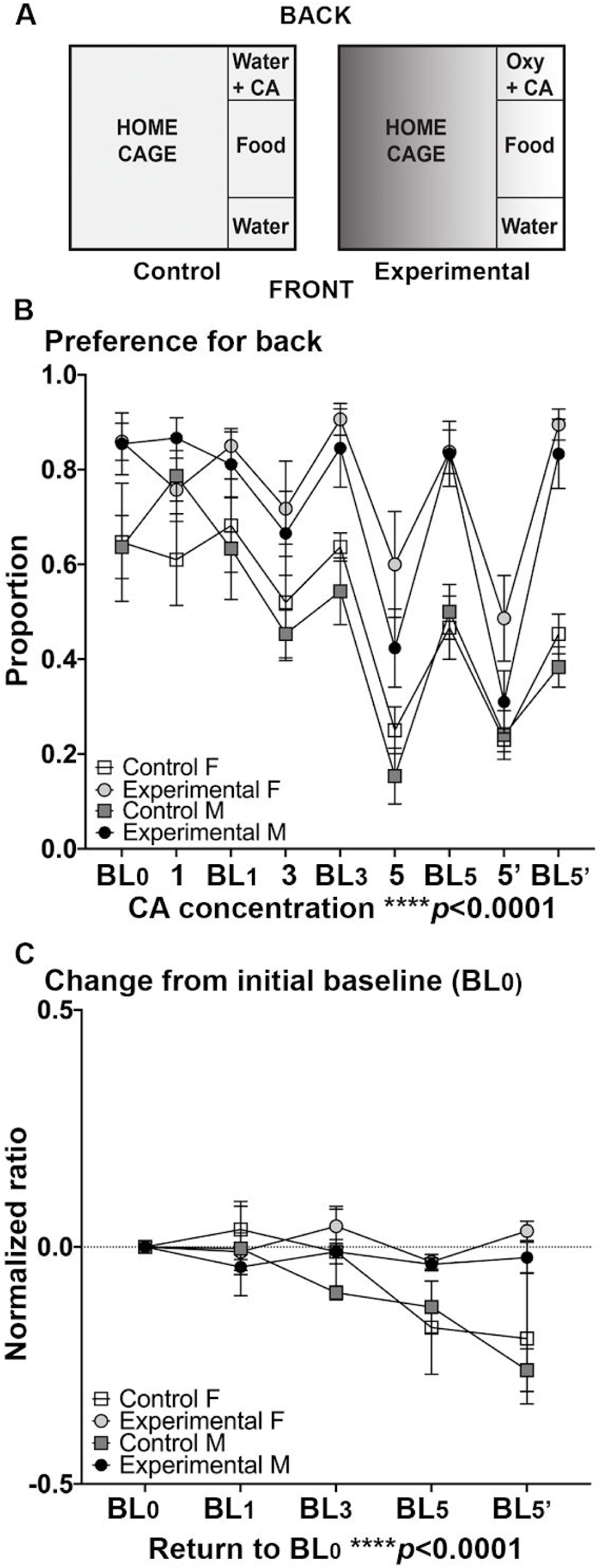
Citric acid suppressed drinking in all rats, but experimental - but not control - rats returned to baseline preference. **(A)** Schematic of home cage layout during the citric acid (**CA**) test. Control rats (left panel) had a water bottle with increasing concentrations of CA every third day (1, 3, and 5 mM) in the back an unadulterated water bottle in the front. Experimental rats (right schematic, shaded) received a water bottle in the front and an oxycodone bottle (**Oxy**) with increasing concentrations of CA every third day (1, 3, and 5 mM) in the back. The 5mM concentration of CA was repeated twice (5 and 5’). CA days were interspersed with “baseline” days (**BL0, BL1, BL3, BL5, BL5’**). **(B)** Preference for the back bottle of Control and Experimental rats analyzed via two-way RM ANOVA (CA concentration x Treatment) revealed a significant interaction F(24,152=1.621, ***p*=0.0432**, significant effect of CA concentration F(8,152)=26.5, ***p*<0.0001**, and a significant effect of Treatment F(3, 19)=4.806, ***p*=0.0118**. Post-hoc analysis of the preference change over increasing concentrations of CA revealed Experimental males but not females suppressed at 3mM CA (***p*=0.0275**) compared to the initial preference when there was no citric acid (CA concentration=0; **BL0**). All animals significantly suppressed intake at both presentations of the 5mM concentration: Experimental females (5, ***p*=0.0018** and 5’ ***p*<0.0001**); Control females (5, ***p*=0.0041** and 5’ ***p*=0.0022**); Control males (5, ***p*=0.0003** and 5’, ***p*=0.0041**); Experimental males (5 and 5’ ***p*<0.0001**). **(C)** Analysis of change from initial baseline (**BL0**) via a two-way RM ANOVA (Return to BL_0_ x Treatment) revealed a significant interaction F(12,76)=3.851, ***p*=0.0001**, an effect of Return to BL_0_ F(4, 76)=9.591, ***p*<0.0001**, and no effect of Treatment F(3,20)=1.784, *p*=0.1825. Post-hoc analyses showed that Control female and Control male rats maintained their aversion compared to **BL0** intake (Control females **BL5, *p*=0.0151**; **BL5’ *p*=0.0046**; Control males **BL5’, *p*<0.0001**), whereas Experimental female and male rats returned to BL0 levels (Experimental females **BL5**, p=0.7879, **BL5’**, p=0.7414; Experimental Males BL5, p=0.6400; **BL5’**, p=.9067).

### 3.4. Pre-screening of open field behavior predicted oxycodone intake-related outcomes in both female and male rats

To provide an unsupervised picture of the relationship of open field behavior to dependent measures during the oxycodone intake phase (intake, preference, blood levels, precipitated withdrawal, spontaneous withdrawal, CA, gnawing, etc.), we conducted a factor analysis of all pre-screening open field behaviors and marble burying. The analysis revealed a single factor for both females and males that explained ∼71% of the variance each. The scatterplots for these analyses are shown separately for females **(Fig. 6A)** and males **(Fig. 6B)**. To obtain a single score for each rat, the pre-screening open field behavior measures were multiplied by the factor score of that behavior and summed. Each rat’s factor score was then correlated with their dependent measures during the oxycodone intake phase. For females, only gnawing was significantly correlated with the open field measure. For males, significant correlations were seen with precipitated withdrawal weight loss, gnawing, oxycodone dose at Week 22, liquid intake at Week 22, and blood levels **(Fig. 6C)**.

**Figure 6.**
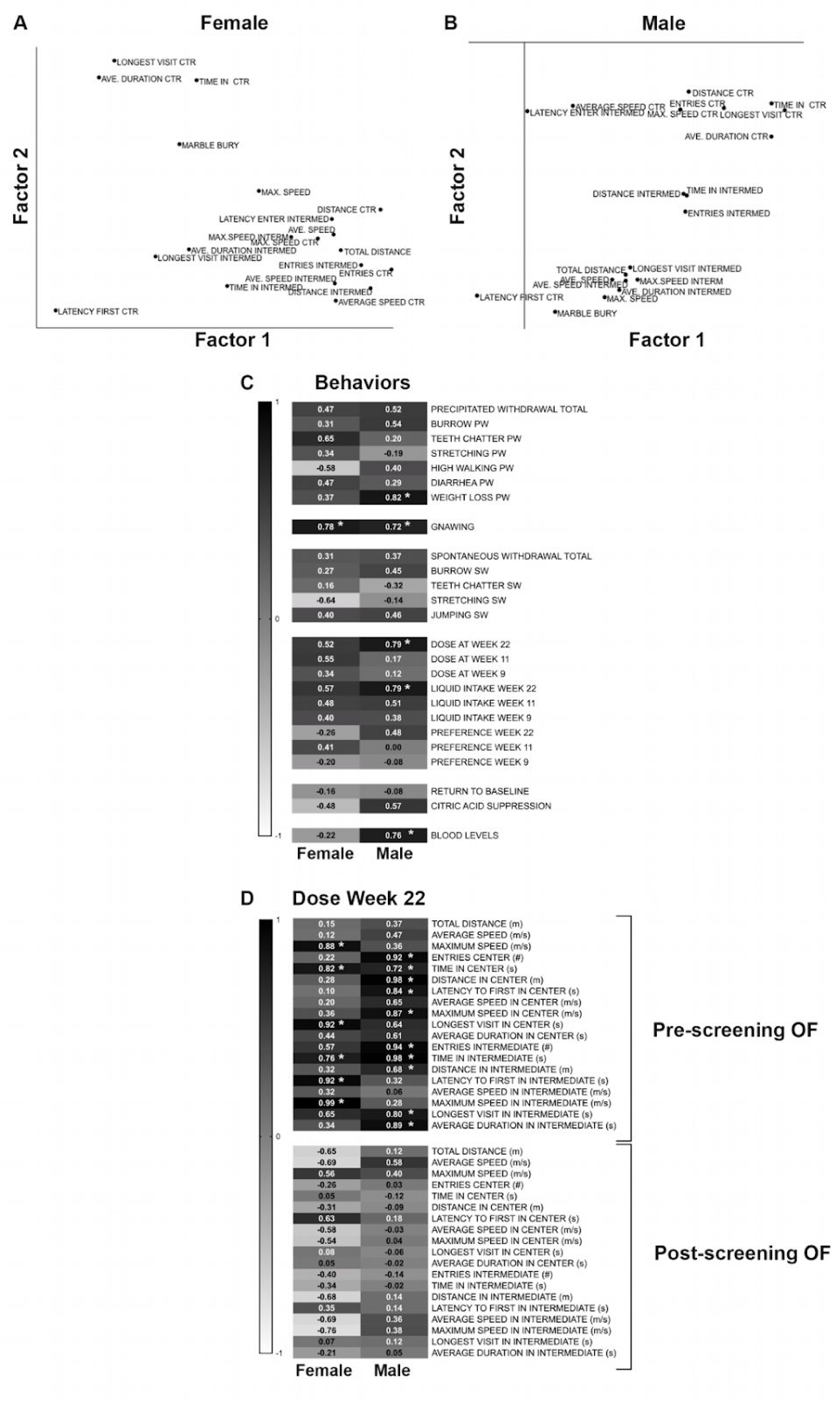
Factor analysis (FA) and Correlations among pre-screening behavior and post-oxycodone outcomes. **(A)** The scatterplot of the factor analysis of the behavioral data for the pre-test open field and marble burying in females. Factor 1 on the x-axis (70.99% of the variance) is plotted against the Factor 2 on the y-axis (9.94% of the variance). Factor 1 loadings from the open field behaviors were used in Panels C and D. **(B)** The scatterplot of the factor analysis of the behavioral data for the pre-test open field and marble burying in females. Factor 1 on the x-axis (70.63% of the variance) is plotted against the Factor 2 on the y-axis (15.57% of the variance). Factor 1 loadings are used in Panels C and D. **(C)** Each behavior was weighted by the loadings on Factor 1 and summed to provide an overall pre-test score. This heat-map presents the multiple correlations (+1.0 to −1.0) between that Factor 1 score and the post-oxycodone variables related to intake and withdrawal. For females only gnawing significantly correlated with the pre-test score (r>0.707; n=8). For males, precipitated weight loss, gnawing, oxycodone dose self-administered at Week 22, liquid intake at Week 22 and oxycodone blood levels significantly correlated with the pre-test score (r>0.666; n=9). **(D)** Heat-map of the multiple correlation between the self-administered dose at Week 22 and the pre- and post-screening open field variables in both females and males. The significant *p-*value is highlighted by the white star and for females is 0.707 (n=8) and 0.666 for males (n=9).

In addition to the factor analysis described above, analysis of the correlation data between pre-screening open field behavior and oxycodone intake at Week 22 revealed a positive relationship between activity measurements in the open field, particularly increased activity in the center and intermediate areas, and intake **(Fig. 6D)**. In contrast, there were no significant correlations between oxycodone intake at Week 22 and post-screening open field (OF2) behavior **(Fig. 6D)**.

## 4. DISCUSSION

Important aspects of a preclinical model for human prescription opioid abuse liability include prolonged access to drug, voluntary and often increasing self-administration when a choice is provided, high levels of intake, drug dependence, and subsequent somatic signs of opioid withdrawal. Here we show that when provided to rats via a classic voluntary drug intake model (2-bottle choice), oxycodone recapitulates these key features of human prescription opioid abuse. Rats given chronic, continuous choice between oxycodone and water reliably demonstrate voluntary and escalating intake of oxycodone, measurable levels of dependence, and motivation to take the drug. We believe this preclinical model of prescription opioid abuse - which relies on oxycodone oral self-administration - is of significant translational relevance for understanding the mechanisms that underlie opioid addiction and ultimately will contribute to the ongoing research efforts to combat the growing opioid epidemic.

### 4.1. Conceptual and methodological pros and cons of long-term, voluntary, oral oxycodone intake via 2-bottle choice

The face validity of opioid self-administration models is based on an animal’s motivation to seek and to take drugs (Spanagel, 2017) and to do so despite adverse consequences (Koob and Le Moal, 2001). Employing a classic 2-bottle choice paradigm, rats readily drank oxycodone and showed a preference for the oxycodone-containing bottle relative to the water bottle. Experimental rats escalated their oral oxycodone intake over time, suggesting it was reinforcing, and rebounded to these high levels of intake following citric acid suppression. Thus, rats in a chronic, continuous oxycodone 2-bottle choice model show motivation to drink in the face of adverse consequences.

Other opioid self-administration paradigms have relied on either oral intermittent access (OIA) or intravenous (IV) self-administration (Carroll and Meisch, 1984; Enga et al., 2016; Gellert and Holtzman, 1978a; Grim et al., 2018; Heyne, 1996; Jimenez et al., 2017; Leung et al., 1986; Meisch, 2001; Nguyen et al., 2018; Nichols and Hsiao, 1967; Shaham et al., 1993; You et al., 2017). Both OIA and IV self-administration models can lead to dependence and tolerance, and IV models can demonstrate drug-seeking motivation. OIA models mimic a common route of administration of humans (oral), whereas the rapid onset of IV drug effects enables assessment of contingent cues and behaviors and mimics IV drug users. However, these models have limitations. High first-pass metabolism of drugs in OIA opioid models results in a short bioavailability and longer latencies to induce the “high” compared to IV models (Chan et al., 2008). IV self-administration models often rely on short-term access, which confounds the assessment of drug effects vs. those caused by daily withdrawal (Badiani et al., 2011; Meisch, 2001; Weise-Kelly and Siegel, 2001). We believe the oxycodone 2-bottle choice model used here addresses some of these issues. Its advantages include: a) the translational relevance for orally-delivered drugs, b) voluntary intake and choice between natural rewards (food/water) and oxycodone, c) the chronicity of drug available 24h/7d over months, d) the non-invasive methodological approach, and e) the absence of confounders, such as adulteration of oxycodone solution to encourage oral intake or stress the animal to induce intake.

In comparison to the multiple studies in which opiates were administered orally but intermittently (Davis et al., 2010; Enga et al., 2016; Fan et al., 2018; Gellert and Holtzman, 1978a; as reviewed in Meisch, 2001), there are fewer studies that employed long-term continuous oral intake of morphine or other opioids. Those more chronic studies started with low opioid concentrations that were increased by the experimenter (Badawy et al., 1982), mixed with a sucrose solution (Fuentes et al., 1978; Leung et al., 1986), or provided as the sole source of liquid (Gellert and Holtzman, 1978a; Shaham et al., 1992), with ingestion often limited to a few weeks. To our knowledge, no studies to date allowed the rodent to freely self-titrate opioid intake over several months without some form of taste enhancement or liquid constraint. In the present study, female and male rats preferred and drank increasing amounts of oxycodone over weeks and months; in prior shorter duration studies, rats did not prefer the opioid (typically drinking < 50% of total liquid intake) and did not escalate their intake (Badawy et al., 1982; Gellert and Holtzman, 1978a), with the exception of the opioid etonitazene (McMillan et al., 1976). When tested, in all studies of oral opioid intake, including the present one, rodents displayed withdrawal behaviors when challenged with an opioid antagonist, highlighting the ability of many different oral paradigms to result in opioid dependence.

One caveat to the present study is that the 2-bottle paradigm used here necessitated that the rats be individually housed for ∼10 months, and thus the results must in interpreted in the context of this social isolation. Social isolation augments many effects of opioids including oral self-administration and locomotor activity (Alexander et al., 1981; Coudereau et al., 1996; Deroche et al., 1994; Francès et al., 2000; Hadaway et al., 1979; Sudakov et al., 2003), but reduces morphine-induced place preference and analgesia (Bozarth et al., 1989; Coudereau et al., 1997; Schenk et al., 1983; Wongwitdecha and Marsden, 1996). If isolation indeed reduces the effects of opioids results in increased compensatory intake, the effects of isolation would be sex-dependent since the preference, intake, and blood levels were elevated in females compared to males. Future work could test whether the high escalating levels of oxycodone drinking we find in females and males and the sex differences in that intake would also occur in socially-housed animals.

### 4.2. Oxycodone 2-bottle choice highlights sex differences in oxycodone intake and blood levels

Prescription opioid abuse liability is increasingly understood as being sex-dependent (Becker and Koob, 2016; Bobzean et al., 2014), which is supported by the more rapid increase in the rate of prescription opioid abuse in women than men (Warner et al., 2016). Previous work suggests a role for sex hormones in opioid dependence and opioid reinforcing properties (Alexander et al., 1978; Bobzean et al., 2014; Cicero et al., 2003, 2002b, 2002c; Hadaway et al., 1979; Harte-Hargrove et al., 2015; Serdarevic et al., 2017), particularly in oral self-administration protocols (Stolerman and Kumar, 1972). Sex differences are also apparent in the pharmacokinetic, metabolic, and analgesic effects of oxycodone (Chan et al., 2008; Holtman and Wala, 2006; Neelakantan et al., 2015), and subtle differences have been observed in the liability for opioid abuse (Collins et al., 2016; Mavrikaki et al., 2017). Therefore animal studies show that, relative to males, females self-administer more opioids, are more vulnerable to their reinforcing effects, and become more physically dependent, (Alexander et al., 1978; Boyer et al., 1998; Cicero et al., 2003, 2002a; Graziani and Nisticò, 2016b; Hadaway et al., 1979; Lofwall et al., 2012; Lynch, 2018; Lynch et al., 2002; Serdarevic et al., 2017). In our work presented here, when rats were given a choice between a water bottle and an oxycodone-containing bottle, both sexes readily drank oxycodone and escalated their intake, but females self-administered twice as much oxycodone by body weight as did males, with a resultant five-fold increase in blood levels. The preference for oxycodone shown by females peaked early and remained at high levels throughout, whereas this peak in preference took longer for males. Notably, these differences were independent of oxycodone concentrations since experimental groups escalated their intake over the course of chronic exposure. Moreover, in the oxycodone 2-bottle choice paradigm presented here, individual differences during pre-screening were less predictive of intake, self-administered dose, blood levels, and gnawing behavior for females than males. Taken together, these data suggest that the chronic, continuous oxycodone 2-bottle paradigm used here is suited to interrogate sex differences under a choice self-administration paradigm.

In humans, females have higher CYP3A4 activity suggesting a faster oxycodone elimination than males, with females displaying more vulnerability to oxycodone’s effects (Andreassen et al., 2011; Graziani and Nisticò, 2016b). Although we did not investigate sex differences in pharmacokinetics, the large difference in dosing observed in our paradigm could have been driven by sex differences in oxycodone metabolism. When oxycodone was administered by gavage to Sprague Dawley rats, females showed higher oxycodone blood levels than males, providing evidence for differences in oxycodone metabolism (Chan et al., 2008). However, other work found subtle differences in IV self-administration but no differences in oxycodone blood and brain levels, suggesting no pharmacokinetic differences (Mavrikaki et al., 2017). It is tempting to speculate that the higher total liquid intake in females in the data presented here was due to a higher intake needed to keep oxycodone blood levels constant. This would argue in favor of a faster elimination rate (but see Chan et al., 2008), lower oxycodone sensitivity and/or different first pass metabolism, resulting in greater intake at the same concentration compared to males. Further studies on blood sampling at earlier and multiple time points along with metabolite measures are warranted to test this hypothesis.

### 4.3. Oxycodone two-bottle choice as a model of dependence and stereotyped gnawing behavior

With the long-term free-access oxycodone intake used, most rats will escalate their drug intake over time, which contrasts with the lack of escalation in many short-term access paradigms (Badiani et al., 2011; Meisch, 2001). In addition, presumably chronic, continuous access allows rats to titrate their intake to avoid withdrawal. In support of this, spontaneous withdrawal signs were not statistically different between control and oxycodone rats, suggesting that spontaneous withdrawal in our long-term access paradigm is negligible. There are caveats though; our sampling for spontaneous withdrawal was time- and behavior-limited, and a more thorough analysis of spontaneous withdrawal signs in the oxycodone 2-bottle choice model is warranted. In contrast to the lack of spontaneous withdrawal, both female and male experimental rats showed robust precipitated withdrawal signs (Bläsig et al., 1973; Gellert and Holtzman, 1978a; Leung et al., 1986; Sudakov, 1991), with ∼4-fold more naloxone-induced signs of physical dependence relative to control rats. Thus, the oxycodone 2-bottle choice model will be useful for assessing opioid dependence and for assessing the efficacy of treatments targeted to relieve withdrawal.

Despite no significant differences in spontaneous withdrawal, in the oxycodone two-bottle choice model used here Experimental (but not Control) rats developed stereotyped gnawing behavior. Gnawing and other oral stereotyped behaviors can be induced by repeated high doses of morphine (Wennemer and Kornetsky, 1999) or by stress and are mediated by endogenous opioids (Bergmann et al., 1974; Ernst and Smelik, 1966; Morelli et al., 1989). One source of stress in the two-bottle choice model is social isolation which reduces tolerance to morphine but also precipitated withdrawal (Broseta et al., 2005; Coudereau et al., 1996; Jiménez and Fuentes, 1993). Here rats were singly-housed for 10 months, longer than in the aforementioned citations; however, isolation is unlikely the sole contributor to increased gnawing behavior as no control rats measurably gnawed. It is possible that the combination of oxycodone and isolation increased the likelihood of gnawing, and we might expect that group-housed oxycodone rats would gnaw less than isolated oxycodone rats. Moreover, because social isolation is more stressful for females than males (Beery and Kaufer, 2015; Hatch et al., 1965; Westenbroek et al., 2005), the increased gnawing in particular by experimental females may be explained by an opioid-mediated activation of the dopaminergic mesolimbic system known to induce gnawing behavior (Bergmann et al., 1974; Morelli et al., 1989).

### 4.4. Oxycodone two-bottle choice: appetitive and reinforcing behavior, escalation, and incentive salience

When challenged to drink an aversive tastant added to the oxycodone bottle, most rats, (Experimental and Control) suppressed their drinking from the citric acid adulterated bottle. However, some Experimental rats maintained their preference for oxycodone despite the addition of citric acid. Both female and male experimental rats readily returned to baseline levels when the unadulterated oxycodone bottle was returned, whereas control rats maintained their acquired aversion. These data suggest that two-bottle choice of oxycodone has high face validity for modeling prescription opioid abuse liability and it provides a broader use of this model to study motivational aspects of prescription opioid abuse, such as individual differences in the development of physical dependence, oxycodone reinforcement, and its motivational salience.

### 4.5. Pre-screening for predictive factors

Not all humans become addicted to drugs and those that do show different degrees of dependence (Brady and Randall, 1999; Cicero and Ellis, 2017b). We believe the individual differences that emerge during the oxycodone two-bottle choice model can be harnessed to identify individual vulnerabilities that may be predictive of oxycodone abuse liability. Previous work defining predictive factors that contribute to an addictive phenotype (Belin et al., 2016) shows that locomotor and exploratory activity and anxiety-like phenotypes can be predictive of later drug-seeking behavior (Markel’ et al., 1989; Piazza et al., 1989). In our present work, there were sex differences in the prescreening prediction of later intake and intake-related variables; pre-screened open field behaviors predicted intake and intake-related variables and some components of dependence in males but not females. A caveat of this conclusion is that females had consistently higher levels of oxycodone intake than males. Thus, the fewer predicted intake specific variables for females compared to males may reflect a ceiling effect specifically for intake variables. This ceiling effect is less likely to explain the lack of correlation for non-intake-related variables such as weight loss after precipitated withdrawal.

### 4.6. Conclusions

Relying on a large literature using two-bottle choice models of drug intake (Hill and Powell, 1976), we show that rats voluntarily drink increasing amounts of oxycodone when given continuous access and prefer it to water when given a choice between oxycodone and water. In addition to mimicking oral ingestion of oxycodone in humans, the two-bottle choice model we use here presents several advantages as an additional model to study the abuse aspects of this drug, including the provision of choice between water and drug and unlimited access to the drug, which allows the animal to titrate its oxycodone levels in contrast to experimenter-controlled dosing. When allowed to freely drink oxycodone, both females and males increase their intake over time, show physical dependence as indicated by precipitated withdrawal and stereotyped gnawing behavior. Oxycodone drinking Experimental rats rebound from an aversive citric acid challenge whereas Control rats do not. Females escalate more rapidly, show higher blood levels, greater levels of dependence and more gnawing, setting the stage for the study of sex differences in the motivation to take oxycodone and the consequences of that intake.

Future work would address why and how individual rats differ in their intake, pre-existing risk factors that might predict high levels of intake, mechanisms underlying the sex differences, and the role of isolation in these studies. We believe that the oxycodone two-bottle choice model is germane to probing treatment options, pharmacological and environmental, and in particular the role of stress and isolation in the motivation to take opioids. Finally, as there are differences in brain structure activation in oxycodone exposed verse naive rats given acute oxycodone (Iriah et al., 2019), imaging rats

pre-and post-oxycodone in females and males during the course of their voluntary intake could provide insights into neurocircuitry of oxycodone use in the two sexes.

## ACKNOWLEDGEMENTS

We thank Ashlee A. Dougher, Peter Lenchur, Patrese

Robinson-Drummer, and Steve J. Simmons with technical assistance during data acquisition, and Ganesh S. Moorthy, Christina M. Vedar, and Athena F. Zuppa of CHOP’s Bioanalytical Core Center for Clinical Pharmacology for oxycodone analysis in blood.

